# Vector competence and transcriptional response of *Aedes aegypti* for Ebinur Lake virus, a newly mosquito-borne orthobunyavirus

**DOI:** 10.1101/2022.02.14.480372

**Authors:** Cihan Yang, Fei Wang, Doudou Huang, Haixia Ma, Lu Zhao, Guilin Zhang, Hailong Li, Qian Han, Dennis Bente, Zhiming Yuan, Han Xia

## Abstract

The global impact of mosquito-borne diseases is increasing in the last decades. The newly classified orthobunyavirus, Ebinur Lake virus (EBIV) has been verified with highly virulent pathogenic to adult laboratory mice, and antibodies against EBIV have been detected in humans. As a potential emerging virus, it is necessary to assess the vector capacity of mosquitoes for EBIV to predicting its risk to public health. Herein, *Aedes aegypti*, the gradually important vector in China, was used as a model to evaluate the vector competence for EBIV. It was showed that EBIV can be transmitted by *Ae. aegypti* through oral feeding and the transmission rates could get to 11.8% at 14 days post infection (dpi). The highest infection rate, dissemination rate and ovary infection rate were 70%, 42.9%, and 29.4%, respectively. Through intrathoracic infection, *Ae. aegypti* was highly susceptible to EBIV and the transmission rates could get to 90% at 10 dpi. Moreover, the infection rate, dissemination rate and ovary infection rate were all 100%. Transcriptome analysis demonstrated EBIV can alter expressions of mosquito genes related to immune-related process and metabolism-related process. *Defensin-C* and *chitinase 10* had been continuously down-regulated in the mosquitoes infected by intrathoracic inoculation. Our studies made the comprehensive analysis of the vector competence and transcriptional response of *Ae. aegypti* for EBIV, which implied the potential risk of EBIV to public health. Moreover, these findings indicated a complex interplay between EBIV and the mosquito immune system to affect the vector transmission capability.

## 1. Introduction

Mosquito-borne viruses (MBVs), as a group of heterogeneous RNA viruses, naturally survive in both mosquitoes and vertebrate hosts, and are the aetiological agents of many human diseases [1]. The medically important MBVs are primarily distributed to three families and one order: the *Flaviviridae* family [dengue viruses 1-4 (DENV), Zika virus (ZIKV), West Nile virus (WNV), Japanese encephalitis virus (JEV), etc], the *Togaviridae* family [Chikungunya virus (CHIKV), etc], the *Reoviridae* order [(Banna virus (BAV), etc) and *Bunyavirales* order [Rift Valley fever virus (RVFV), etc] [2,3]. In the past seven decades, the extensive global spreads of MBVs in the world have been remarkable. For example, the most notable flavivirus DENV, the pathogen of dengue fever, has increased dramatically within the past 20 years, and more than 3.9 billion people in 128 countries are reported to be at risk of dengue infection [4]. After ZIKV and WNV Zika virus are introduced into the Western hemisphere, the viruses have a rapid geographical spread, and cause a large number of infections in the population [5,6]. The dramatic emergence and spread of epidemic mosquito-borne diseases have made it necessary to do surveillance and risk assessment.

The genus *Orthobunyavirus*, belongs to family *Peribunyaviridae* in the order *Bunyavirales*, containing the many mosquito-borne bunyaviruses [7]. So far, the orthobunyaviruses can be discovered in mosquitoes of the different species, such as *Ochlerotatus* spp., *Culex* spp. and *Aedes* spp. [8–10]. Some viruses in this genus can cause severe human diseases, including acute but self-limiting febrile illness (Oropouche virus, OROV), encephalitis (La Crosse virus, LACV) and haemorrhagic fever (Ngari virus, NRIV) [11]. Reports have found that some orthobunyaviruses have a wide epidemic distribution in the population, such as Bunyamwera virus (BUNV) which is endemic in many African countries and North America, and Batai virus (BATV) which is geographically widespread in Europe [12]. With the worldwide spread of medically important mosquitoes, these viruses are also possibly widely spread and pose a potential threat to human public health, animal health, and food security.

*Ae. aegypti* is an important vector of DENV, ZIKV, CHIKV and yellow fever virus (YFV) [13]. For a long time, *Ae. aegypti* was only distributed in the African continent and Southeast Asia, but at present, they have colonized almost all continents [14]. The distribution of *Ae. aegypti* has a continuous growth, and it is estimated that a further increase will be approximately 20% to the end of this century [15]. In China, as a result of warmer daily temperatures from 2004 to 2019, the vector competence of *Ae. aegypti* for DENV has an obvious increase [16]. The status of *Ae. aegypti* as the pathogenic vector has a significant rise interiorly. To address these challenges, it is necessary to comprehensively assess the vector capacity of *Ae. aegypti* to transmit these viruses.

Ebinur Lake virus (EBIV), a newly identified orthobunyavirus in China, is isolated from *Culex modestus* mosquito pools in Xinjiang Province [17]. The whole genome sequence of EBIV shares the highest similarity with Germiston virus only found in South Africa, and viral genomic S and M segment showed 95.0% and 86.1% amino acids similarities, respectively [18]. EBIV can efficiently infect cells derived from rodent, avian, non-human primate, mosquito and human [19]. After the EBIV infection, BALB/c mice show the encephalopathy, hepatic damage and immunological system damages with high mortality [20]. Even though there is no report about confirmed human cases of EBIV, the serological proofs for EBIV infection has been detected [19].

Due to the health risk of EBIV, it is necessary to assess the vector competence of mosquitoes for EBIV. Previous studies on vector competence of mosquitoes orthobunyaviruses have shown that *Ae. aegypti* is a potential vector for BUNV, NRIV and LACV [21–23]. Based on these, we selected *Ae. aegypti* as the vector to evaluate the EBIV infection, dissemination, and transmission in mosquitoes. In addition, the transcriptional response of the mosquitoes elicited by EBIV was also investigated. Though two methods, blood-feeding and intrathoracic injection, we thoroughly studied the vector competence and transcriptional response of *Ae. aegypti* for EBIV, which is beneficial to better prepare for and respond to potential outbreak of EBIV, and to understand the transmission mechanism of orthobunyavriuses.

## 2. Materials and methods

### 2.1 Viruses and cell lines

BHK cells were maintained in Dulbecco’s modified Eagle’s medium (Gibco, United States) containing 10% fetal bovine serum (Bio-one, China) at 37°C with 5% CO_2_. *Aedes albopictus* C6/ 36 cells were grown in Roswell Park Memorial Institute (RPMI) 1640 medium (Gibco, United States) supplemented with 10% FBS (Gibco, United States) at 28°C with 5% CO_2_. All the cells have been routinely confirmed to be mycoplasma free.

The EBIV isolate Cu20-XJ is obtained from *Cx. modestus* mosquitoes in 2013 in Xinjiang, China, by inoculation in suckling mice and passaging four times in BHK-21 cells for the blood meal. EBIV was passaged in C6/36 cells for mosquito infection by microinjection. The viral titers were determined by a plaque formation assay.

### 2.2 Mosquito rearing

Eggs of *Ae. aegypti* (Rockefeller strain) was obtained from Laboratory of Tropical Veterinary Medicine and Vector Biology at Hainan university. The eggs and larvae of *Ae. aegypti* were maintained in a mosquito room with the following condition: 28 °C environment with a light: dark cycle of 12:12 and a relative humidity of 75 ± 5% humidity, approximately. After egg hatching, the larvae were maintained in a shallow dish containing dechlorinate water and ground mouse feed powder. The food was refreshed daily and the water was changed every second day. After five to seven days, the collected pupae were placed in the plastic cups with dechlorinate water. Then put the cups into the mesh cages (30 x 30 x 30 cm), which were kept in the insect incubator with the condition: 28 ± 1 °C environment with a relative humidity of 80% and a light: dark cycle of 12:12. Adult mosquitoes were maintained on 8% glucose solution. Note that mosquitoes were reared in an arthropod containment level (ACL)1 laboratory.

### 2.3 Oral infection and Intrathoracic inoculation

Before oral-infection, five to eight days old, adult mosquitoes were collected by using the mosquito absorbing machine and placed into plastic cups (24oz). And wrap the cup with a cut mosquito net mesh and then cover it with a lid with a hole in the middle. Before the oral-infection and intrathoracic inoculation, female mosquitoes, 5 to 8 d-old in the cups were starved for 24 h.

For oral-infection experiment, the mixture (the defibrated horse blood: supernatant of EBIV-infected BHK cells = 1: 1) were used to feed female *Ae. aegypti* through the artificial mosquito feeding system (Hemotek, UK). Fully engorged female mosquitoes were transferred into new containers and maintained in the incubator at 28 °C in the humidity of 80% for 4 days to 14 days.

For intrathoracic inoculation experiment, anesthetize female mosquitoes at −20°C for one minute and pour them into a petri dish placed on ice and cover the lid quickly. Pick out female mosquitoes with tweezers and put them on the ice plate. Under the dissecting microscope, insert the loaded needle (filled with supernatant of EBIV-infected BHK cells) into the mosquito thorax by using a Nanoject II auto-nanoliter injector (Drummond). Then press the INJECT button, and each mosquito was injected with 100 nl of virus. Injected mosquitoes were transferred into new containers and maintained in the incubator at 28 °C in the humidity of 80% for 2 days to 14 days.

For the estimation of vector competence, three parameters were calculated: infection rate (%) = 100 x (the number of mosquitoes with virus-positive bodies or midguts/ the number of total mosquitoes), dissemination rate (%) = 100 x (the number of mosquitoes with virus-positive heads/ the number of mosquitoes with virus-positive bodies or midguts) and transmission rate (%) = 100 x (the number of mosquitoes with virus-positive saliva / the number of mosquitoes with virus-positive bodies or midguts). The ovary infection rate (%) = 100 x (the number of mosquitoes with virus-positive ovaries/ the number of mosquitoes with virus-positive midguts).

### 2.4 Viral RNA detection by qRT-PCR

For the virus detection of whole mosquitoes, each female mosquito was placed in 300 μl RPMI 1640 medium. For the virus detection of saliva, guts and ovaries, each tissue was dissected and placed into an individual tube with 200 μl RPMI 1640 medium. Each mosquito saliva was collected through the previous protocol [24]. The wings and legs of mosquito were cut off and the mouthpart were inserted into the pipette tips filled with immersion oil. Mosquitoes will secrete saliva into the oil and the process will last for 45-60 min at room temperature. The pipette tip containing the oil and saliva was centrifuged at 5, 000 xg for 5 min at 4 °C to expel saliva into a 1.5 ml tube containing 200 μl RPMI 1640 medium.

Homogenize the above samples by Low Temperature Tissue Homogenizer Grinding Machine (Servicebio, China). Then centrifuge the samples at 12, 000 x g for 10 min at 4°C. Following the instruction, total RNA of each sample was extracted by automated nucleic acid extraction system (NanoMagBio, China). Use the One Step TB Green^®^ PrimeScript^™^ PLUS RT-PCR Kit (Takara) to quantify the viral RNA copies through the CFX96 Touch Real-Time PCR Detection System (Bio-Rad). The primers used for these analyses were shown in Table S1. The cutoff value used for EBIV-positive by qRT-PCR was Ct < 35. The positive cutoff value was determined by comparing paired serial tenfold dilutions either inoculated on cells or assayed by qRT-PCR (Table S2) [25]. The equation of *y* = −3.5566*x* + 37.887 [*x* =lg (the titer of EBIV), *y* = Ct value, R^2^ = 0.9979] was used to calculate the titer of EBIV in each sample.

### 2.5 EBIV antigen detection by immunohistochemistry

The midguts and ovaries were dissected from orally infected female mosquitoes at 14 dpi and intrathoracic-infected female mosquitoes at 7 dpi, respectively. The tissues were fixed by 4% paraformaldehyde for one hour, and then washed with PBST (PBS containing 0.3% Triton X-100) for five times. Next, these tissues were placed in blocking solution (PBS containing 5% goat serum and 0.3% Triton X-100) for one hour. The primary antibody was mice anti-EBIV-NP (diluted 1: 200 in PBST containing 5% goat serum) which was incubated for 24 hours. The secondary antibody was Alexa Fluor 549-conjugated goat anti-mouse IgG (H+L) (diluted 1:250 in PBST containing 5% goat serum, Invitrogen, USA) which was incubated overnight. The actin cytoskeleton was stained by Alexa Fluor™ 594 Phalloidin (Invitrogen) for one hour. After incubation at each step, the tissues should be washed at least five times in 0.3% PBST to avoid affecting following operations. Finally, the tissues were made into slide using SlowFade™ Diamond Antifade Mountants (Invitrogen). Images were recorded with a Leica SP8 confocal microscope (Leica, Germany). Through LAS X software (Leica, Germany), figures were merged and scale bars were added. PowerPoint 2019 was used for image grouping. All samples were analyzed with the same microscope and software settings.

### 2.6 Visualization of EBIV particles by transmission electron microscopy

Ovaries and midguts were dissected from the female mosquitoes infected by intrathoracic inoculation at 7 dpi using a dissecting microscope. These tissues were fixed in 2.5% glutaraldehyde until use. The fixed samples were handled in the Center for Instrumental Analysis and Metrology (Wuhan Institute of Virology, China). They were sectioned using an ultramicrotome, and observed using a Tecnai G20 TWIN transmission electron microscope (FEI, United States). PowerPoint 2019 was used for image grouping.

### 2.7 Transcriptomes analysis

The female mosquitoes infected by intrathoracic inoculation and the mock-infected mosquitoes were collected at 2 and 7 dpi. Ten mosquitoes as a pool were used to extract total RNA using TRIzol reagent (Invitrogen). Three independent biological replicates were performed. The samples were delivered to the Wuhan Benagen Tech Solutions Company for commercial RNA-seq services and data analysis. RNA degradation and contamination were monitored on 1% agarose gels. RNA purity was checked using the NanoPhotometer^®^ spectrophotometer (IMPLEN, CA, USA). RNA integrity was assessed using the RNA Nano 6000 Assay Kit of the Bioanalyzer 2100 system (Agilent Technologies, CA, USA). Sequencing libraries were generated using NEBNext^®^ UltraTM RNA Library Prep Kit for Illumina^®^ (NEB, USA) following manufacturer’s recommendations and index codes were added to attribute sequences to each sample. The clustering of the index-coded samples was performed on a cBot Cluster Generation System using TruSeq PE Cluster Kit v3-cBot-HS (Illumia) according to the manufacturer’s instructions. After cluster generation, the library preparations were sequenced on an Illumina Novaseq platform and 150 bp paired-end reads were generated. Clean reads were obtained by removing reads containing adapter, reads containing ploy-N and low-quality reads from raw data. The clean reads were performed de novo assembly using Trinity program (http://trinityrnaseq.sourceforge.net/). Clean reads were mapped to the *Ae. aegypti* genome database (RefSeq: GCF_002204515.2). The unigene sequences of samples were searched using BLASTX against the Nr, KEGG, GO databases (E-value ≤ 1E-5) to retrieve protein functional annotations based on sequence similarity. The FPKM values were directly used to compare gene expression differences between different samples. The DESeq package was used to obtain the “base mean” value for identifying differentially-expressed genes (DEGs). The absolute value of log_2_ ratio ≥ 1 and *p* value ≤ 0.01 were set as the thresholds for the significance of the gene expression difference between the two samples. Scatter diagrams and bubble diagrams were drawn by GraphPad Prism statistical software 9.1.0.

### 2.8 Data analysis

All data were analyzed by GraphPad Prism statistical software 9.1.0. Differences in continuous variables and differences in mosquito infection rates, dissemination rates, transmission rates and ovary infection rates were analyzed using the non-parametric Kruskal-Wallis analysis for multiple comparisons and Fisher’s exact test where appropriate as indicated in the figure legends, respectively. *p* value ≤ 0.05 was considered statistically significant.

## 3. Results

### 3.1 EBIV can be transmitted by *Ae. aegypti* through oral feeding

In our previous study, when used 10 PFU of EBIV to infect adult female BALB/c mice, the viremia mice can reach to 10^6^ PFU ml^−1^ at 2 dpi [20]. Based on this result, *Ae. aegypti* mosquitoes were fed with a mixture of horse blood and virus supernatant, in which the viral final dose was 3.7 × 10^6^ PFU ml^−1^. The EBIV RNA in these mosquitoes at 4, 7, 10 and 14 dpi were analyzed by qRT-PCR. The mean viral titers of EBIV-positive mosquitoes at the four time points had no significant differences and all of them were above 10^2^ PFU ml^−1^. It was worth noting that some infected mosquitoes were remarkably higher than others and the highest viral titer of these infected mosquitoes could be up to 10^6.4^ PFU ml^−1^ (Fig. 1A). The infection rate at four time points were ranging from 40.3% to 70.7%. The infection rate at 4 dpi was significantly higher than that at 7 (*p* = 0.0086) and 14 dpi (*p* = 0.0005) respectively, and similar with that at 10 dpi (Fig. 1B).

**Figure 1.**
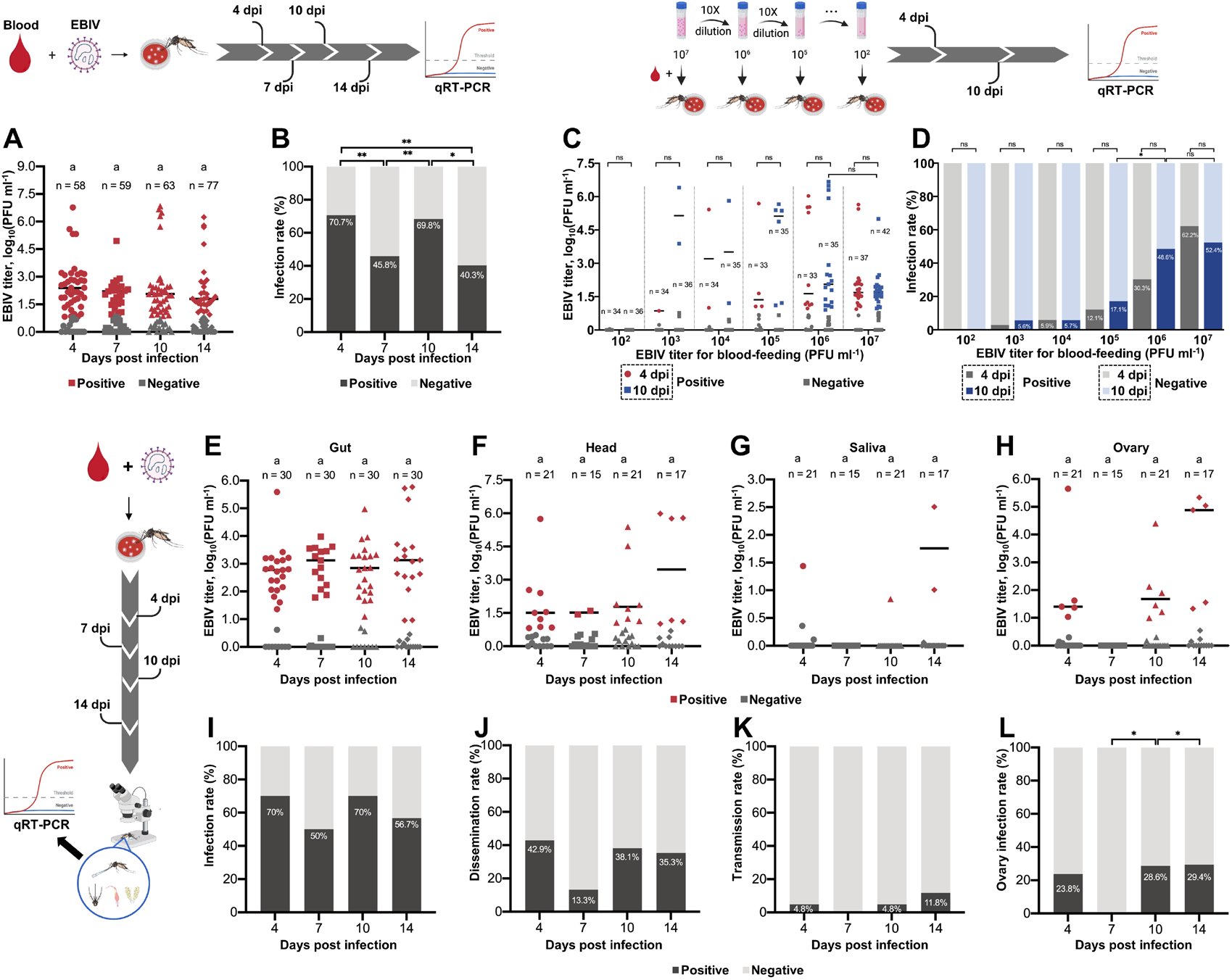
Vector competence of *Ae. aegypti* for EBIV through oral feeding. The EBIV titers (A) and infection rates (B) of mosquitoes at 4, 7, 10 and 14 days after feeding on blood-meal containing 3.7 × 10^6^ PFU ml^−1^ EBIV. The EBIV titers (C) and infection rates (D) of mosquitoes at 4 and 10 days after feeding with six serial viral titers. The EBIV titers in guts (E), heads (F), saliva (G) and ovaries (H) of mosquitoes at 4, 7, 10 and 14 days after feeding on blood-meal containing 6.4 × 10^6^ PFU ml^−1^ EBIV. The infection rates (I), dissemination rates (J), transmission rates (K) and ovary infection rates (L) of mosquitoes at 4, 7, 10 and 14 days after feeding on blood-meal containing 6.4 × 10^6^ PFU ml^−1^ EBIV. Each dot represented an individual mosquito. The same letters are not significantly different (multiple comparisons using the non-parametric Kruskal-Wallis analysis). Differences in the rates were analyzed by by Fisher’s exact test (*: *p* ≤ 0.05, **:*p* ≤ 0.01, ***:*p* ≤ 0.005 and ****:*p* ≤ 0.001).

To confirm the appropriate titer of EBIV for blood feeding, female mosquitoes were fed with six serial viral titers. The EBIV RNA in these mosquitoes at 4 and 10 dpi were analyzed by qRT-PCR. With the increase of EBIV in blood meals, the proportion of infected mosquitoes was gradually rise and the mean viral titers showed no significant difference between 4 dpi and 10 dpi mosquitoes fed with same titer of EBIV (Fig. 1C). The infection rates of the mosquitoes fed with at 10^6^ and 10^7^ PFU ml^−1^ at 4 dpi (30.3%, 62.2%) and 10 dpi (48.6%, 52.4%) were significantly higher than the others, but there were no significant differences between them (Fig. 1D).

In order to examine EBIV distribution in the infected mosquitoes via artificial blood feeding (the viral final titer was 6.4 × 10^6^ PFU ml^−1^), viral RNAs in guts, heads, saliva and ovaries of the mosquitoes at 4, 7, 10 and 14 dpi were determined. The mean titers in the EBIV-positive guts at 4 (10^2.7^ PFU ml^−1^), 7 (10^2.9^ PFU ml^−1^), and 10 (10^2.8^ PFU ml^−1^) dpi were higher than those in heads, saliva and ovaries (Fig. 1E). At 14 dpi the viral titer in heads (10^3.5^ PFU ml^−1^) was similar with the guts (10^3.2^ PFU ml^−1^) at the same point (Fig. 1F). It was worth noting that the viral titer in saliva could get a highest virus titer at 14 dpi (10^2.5^ PFU ml^−1^) (Fig. 1G) and at this time point ovaries also get the highest mean virus titer (10^3.6^ PFU ml^−1^) (Fig. 1H). The infection rates at the four time points were ranging from 50-70%, and it showed no significant difference between them (Fig. 1I). The dissemination rates at the four time points were ranging from 13.3-42.9% (Fig.1 J). The transmission rates at the four time points were ranging from 0-11.8% and the rate was highest (11.8%) at 14 dpi (Fig.1 K). The ovary infection rates at 4, 10 and 14 dpi were 23.8%, 28.6% and 29.4%, respectively. All the ovaries at 7 dpi did not have detectable the virus (Fig. 1L).

### 3.2 EBIV was highly susceptible in *Ae. aegypti* through intrathoracic inoculation

To analyze the affection of the different barriers of mosquitoes on the EBIV transmission, the experiment of virus intrathoracic inoculation was performed. After an injection of 340 PFU EBIV, the mean viral titers of EBIV-positive mosquitoes at 2 dpi (10^5.5^ PFU ml^−1^, H=48.01, *p* < 0.0001) were significantly higher than those at 4, 7 and 10 dpi. The mean viral titer at 14 dpi (10^5.3^ PFU ml^−1^) was the second highest at five time points (Fig. 2A). All of the infection rates at these five time points were 100% (Fig. 2B).

**Figure 2.**
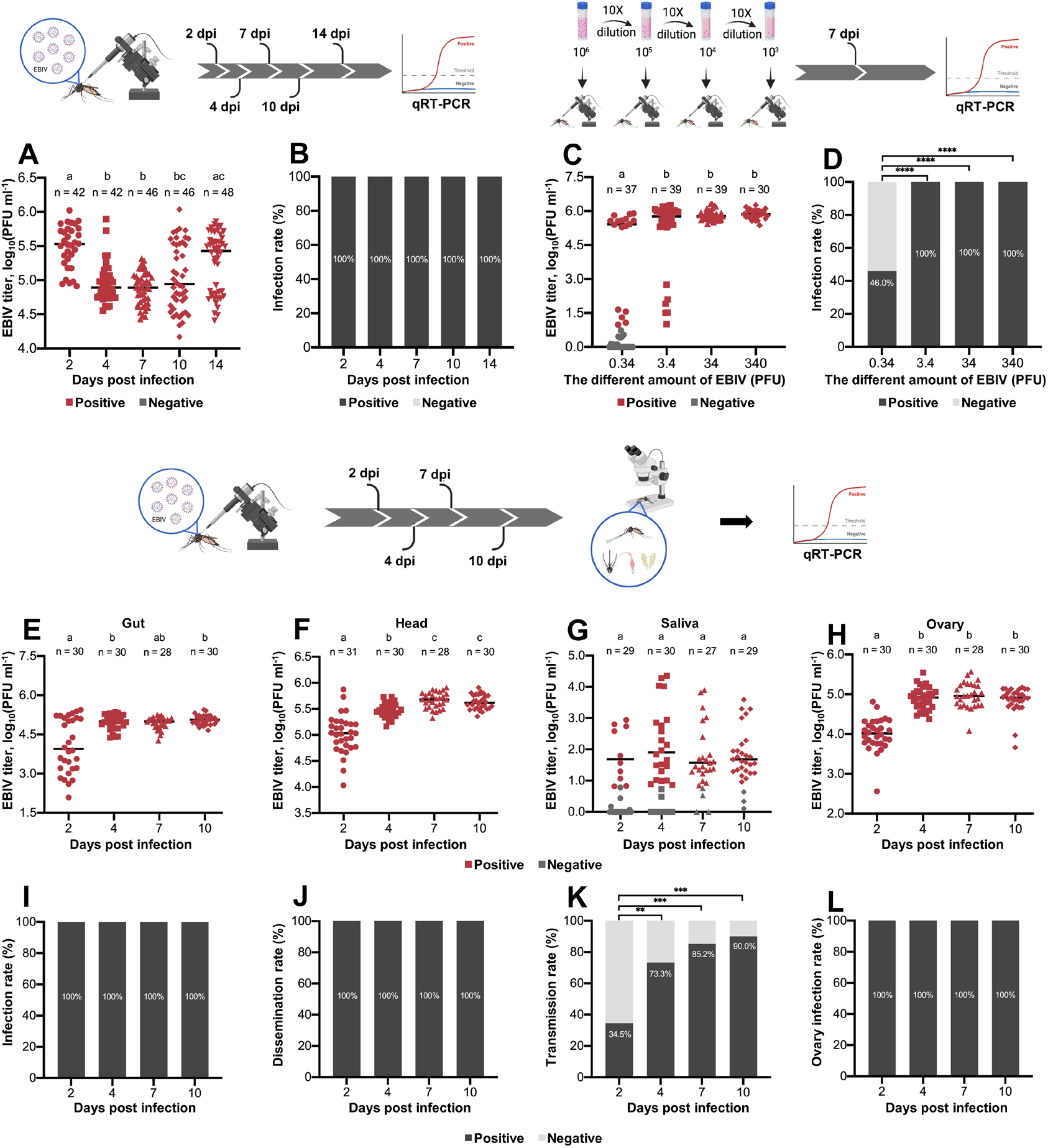
Vector competence of *Ae. aegypti* for EBIV through intrathoracic inoculation. The EBIV titers (A) and infection rates (B) of mosquitoes at 4, 7, 10 and 14 days after injected with 340 PFU EBIV. The EBIV titers (C) and infection rates (D) of mosquitoes injected with four serial viral titers at 7 dpi. The EBIV titers in guts (E), heads (F), saliva (G) and ovaries (H) of mosquitoes injected with 34 PFU EBIV at 2,4, 7 and 10 dpi. The infection rates (I), dissemination rates (J), transmission rates (K) and ovary infection rates(L) of mosquitoes injected with 34 PFU EBIV at 2,4, 7 and 10 dpi. Each dot represented an individual mosquito. The same letters are not significantly different (multiple comparisons using the non-parametric Kruskal-Wallis analysis). Differences in the rates were analyzed by Fisher’s exact test (*: *p* ≤ 0.05, **: *p* ≤ 0.01, ***: *p* ≤ 0.005 and ****:*p* ≤ 0.001).

In order to confirm the appropriate titer of EBIV for the intrathoracic inoculation, female mosquitoes were injected with four serial viral titers. The EBIV RNA in these mosquitoes at 7 dpi were detected by qRT-PCR. The mean viral titer of EBIV-positive mosquitoes injected with 0.34 PFU EBIV was the lowest (10^4.3^ PFU ml^−1^). The mean viral titer of EBIV-positive mosquitoes injected with 3.4, 34 and 340 PFU EBIV were 10^5.1^ PFU ml^−1^, 10^5.8^ PFU ml^−1^ and 10^5.9^ PFU ml^−1^, respectively and there were no significant differences between them (Fig. 2C). The infection rates of these mosquitoes were 100%, except that injected with 0.34 PFU EBIV, having the infection rate of only 46% (Fig. 2D).

### 3.3 The gut of *Ae. aegypti* was the main barrier for the EBIV transmission

In order to examine EBIV distribution in the mosquitoes infected by intrathoracic inoculation (the viral final dose was 34 PFU per mosquito), viral RNAs in guts, heads, saliva and ovaries of mosquitoes at 2, 4, 7 and 10 dpi were determined. The mean titers in the EBIV-positive guts at 4 dpi (10^5.0^ PFU ml^−1^) and 10 dpi (10^5.1^ PFU ml^−1^) were significantly higher than that at 2 dpi (10^4.1^ PFU ml^−1^, H = 13.70 and *p* = 0.0033) (Fig. 1E). The mean titers in the EBIV-positive heads were gradually increased at the four time points and the highest value was 10^5.7^ PFU ml^−1^ at 7 dpi (H = 63.88 and *p* < 0.0001) (Fig. 1F). Notably, the viral titer in saliva at the four time points had no significant difference and the highest value could get to 10^2.2^ PFU ml^−1^ at 4 dpi (H = 20.27 and *p* = 0.0001) (Fig. 1G). And the viral titers in saliva and ovaries also could get very high values at 4 dpi (10^4.9^ PFU ml^−1^), 7 dpi (10^5.0^ PFU ml^−1^) and 10 dpi (10^4.9^ PFU ml^−1^) except that at 2 dpi (10^4.0^ PFU ml^−1^) (Fig.1H). The infection rates, dissemination rates and ovary infection rates were 100 % (Fig. 1I, J and L). With the increase of days after infection, the transmission rates were also slightly increased, ranging from 34.5-90%. The highest transmission rate was at 10 dpi, up to 90% (*p* < 0.0001, the comparison between 2 dpi −10 dpi) (Fig.1 K).

### 3.4 EBIV antigen was detected in the infected mosquitoes

Immunohistochemistry revealed EBIV antigen in midguts of oral-infected mosquitoes at 14 dpi (Fig. 3A), and in midguts and ovaries of the mosquitoes infected by intrathoracic inoculation at 7 dpi (Fig. 3B and 3C). EBIV antigen had accumulated around cytoskeleton in midgut endothelial cells (Fig. 3A and 3B). In the ovarioles, the viral signals were accumulated around nucleus in nurse cells, not in follicle cells (Fig. 3C). EBIV antigen could not be detected in tissues from uninfected mosquitoes (Fig. 3D–3F).

**Figure 3.**
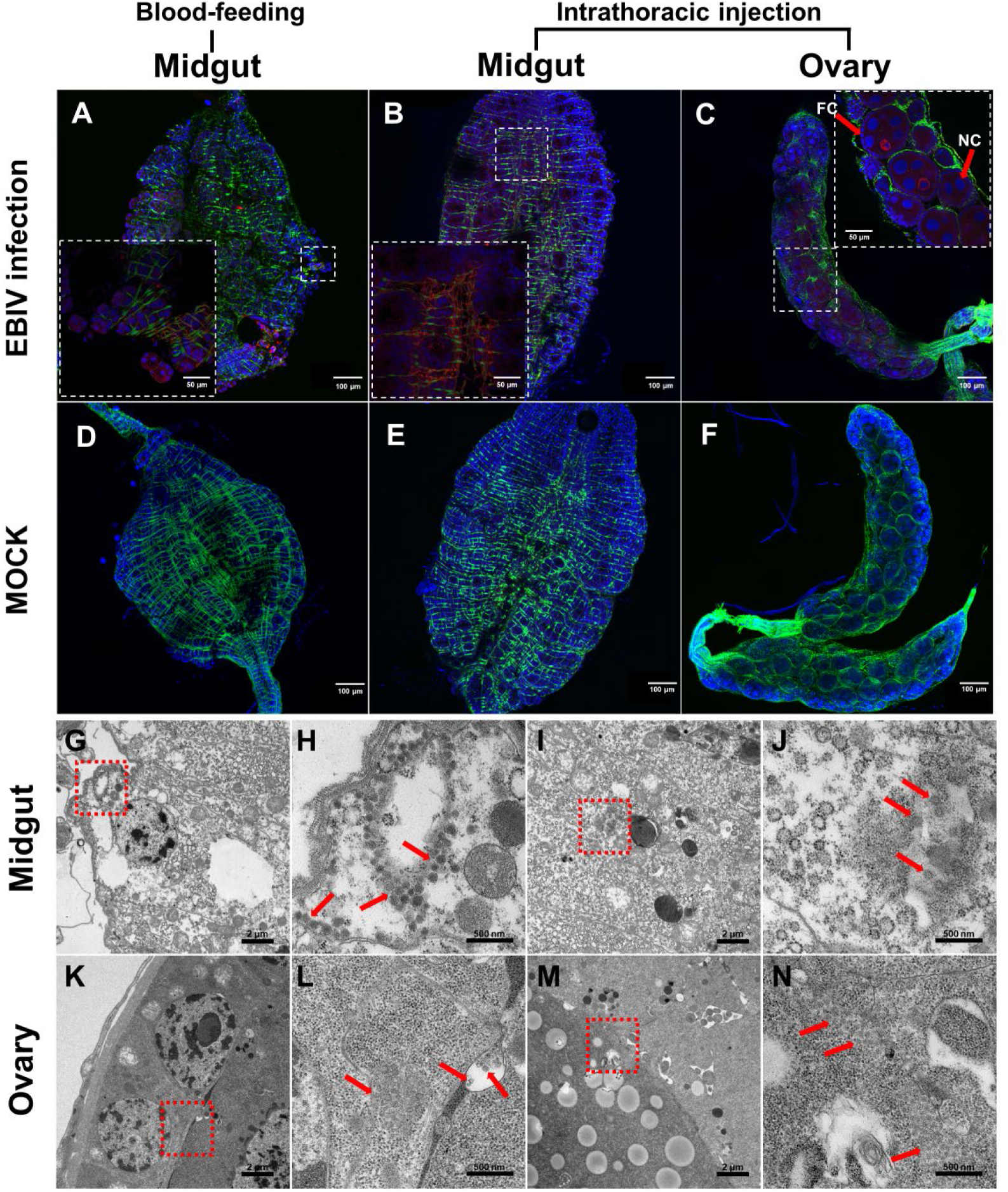
Immunohistochemical visualization and electron micrographs of EBIV antigen and particles in *Ae. aegypti* midguts and ovaries. Immunolocalization of EBIV antigen in midguts from the oral-infected mosquitoes at 14 dpi (A) and from the mosquitoes infected by intrathoracic inoculation at 7 dpi (B). Immunolocalization of EBIV antigen in ovaries from the mosquitoes infected by intrathoracic inoculation at 7 dpi (C). Immunolocalization of EBIV antigen in midguts from mosquitoes fed with blood without EBIV at 14 dpi (D) and from mosquitoes injected with 100 nl RPMI 1640 medium at 7 dpi (E). Immunolocalization of EBIV antigen in ovaries from the mock-injected mosquitoes at 7 dpi (F). F-actin was stained with phalloidin (green). Cell nucleus was stained with DAPI (blue). NC: nurse cells, FC: follicle cells. EBIV virion clusters were detected by use of a mouse anti-EBIV polyclonal antibody and goat anti-mouse IgG labeled with red-fluorescent secondary antibody. (G-J) Virions observed in guts as indicated by red arrows on electron micrographs. H and J were the enlarged insets of the box in G and I, respectively. (K-N) Virions observed in ovaries as indicated by red arrows on electron micrographs. L and N were the enlarged inset of the box in K and M, respectively.

### 3.5 EBIV particles had been observed in the infected mosquitoes

To confirm the EBIV particle morphology in the infected mosquitoes, sections of the digestive tract and ovaries from the mosquitoes infected by intrathoracic inoculation were prepared and examined by transmission electron microscopy (TEM). Spherically shaped virus-like particles (VLPs) (78-140 nm in diameter) were stacked in intracellular vesicles and were very obvious in cells of the digestive tract (Fig. 3G–3J). Similarly, VLPs (60-100 nm in diameter) were present in nurse cells of the ovaries (Fig. 3K–3N). VLPs were not found in uninfected mosquitoes.

### 3.6 Several immune-related genes were identified in EBIV-infected *Ae. aegypti*

To understand the molecular process between EBIV and its vector *Ae. aegypti*, we compared the differences of gene expression between the intrathoracic inoculation and mock-injected mosquitoes at the same time point. Transcriptome analysis results showed that the mosquitoes infected by intrathoracic inoculation at 2 dpi led to significant upregulation of 17 genes and downregulation of 28 genes, while significantly increased expression of 45 and decreased expression of 65 genes were seen in the mosquitoes infected by intrathoracic inoculation at 7 dpi (*p* value ≤ 0.05 and |log2foldchange| ≥ 1) (Fig. 4A and 4B).

**Figure 4.**
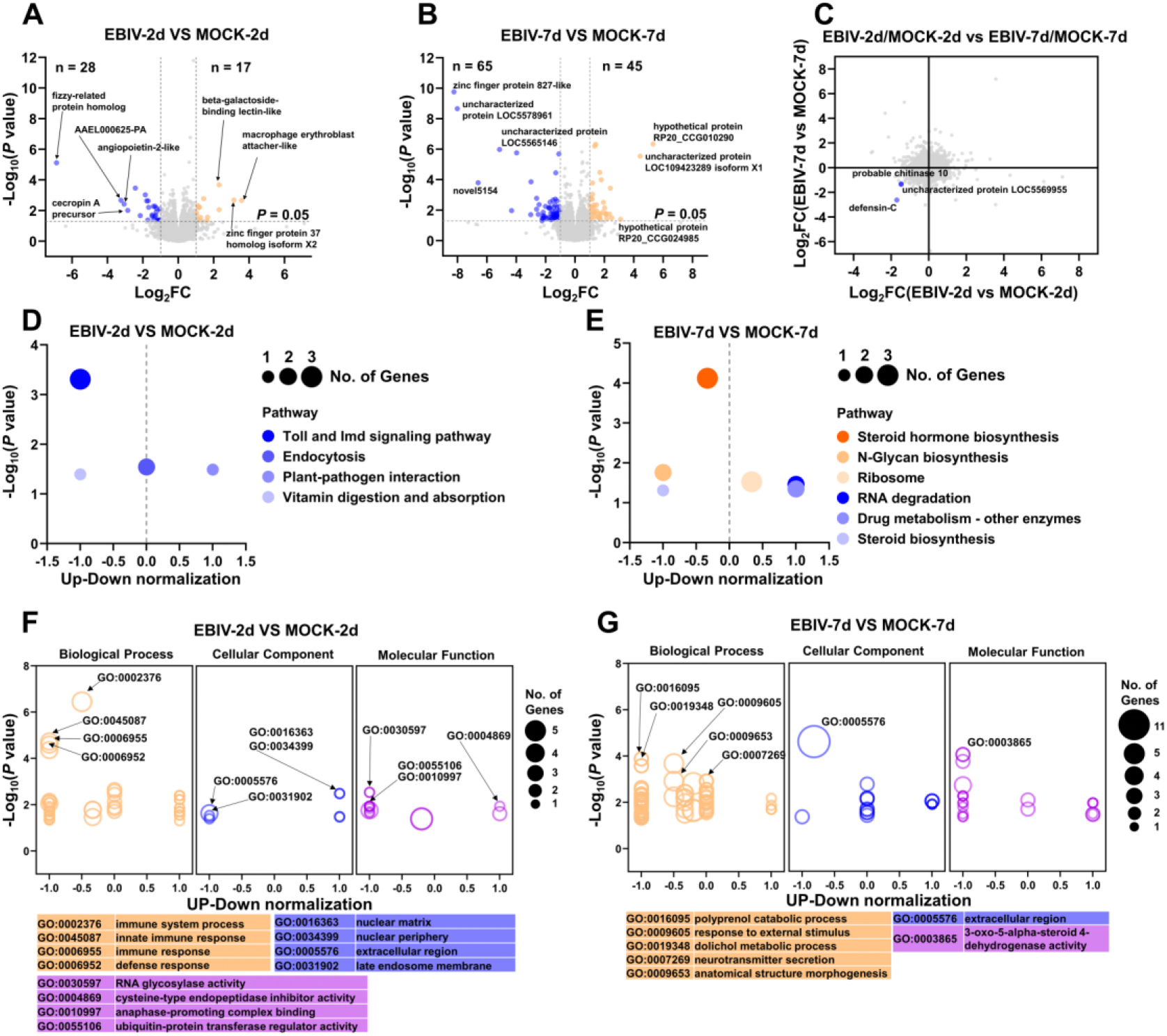
EBIV impacted gene expression in the mosquitoes infected by intrathoracic inoculation. Significantly upregulated (orange) and downregulated (blue) mosquito genes upon EBIV-infected mosquitoes compared with mock-infected mosquitoes at 2 dpi (A) and 7 dpi (B). (C) Genes exhibiting different response to EBIV-infected mosquitoes at 2 dpi and 7 dpi were shown on a scatter plot of expression changes. KEGG pathways analysis of DEGs at 2 dpi and 7 dpi were shown in (D) and (E), respectively. Enriched GO categories associated with DEGs at 2 dpi and 7 dpi were shown in (F) and (G), respectively.

The up-regulated genes were not many in the mosquitoes infected by intrathoracic inoculation at 2 dpi, and most of them were undefined genes (Table S3). The top three proteins expressed by up-regulated genes were macrophage erythroblast attacher-like (E3 Ubiquitin Ligase) related with autophagy [26], zinc finger protein 37 homolog isoform X1/X2 (a transcription factor) and beta-galactoside-binding lectin-like involved the process of virus infection [27] (Fig. 4A). Among the up-regulated genes at this time point, a cuticular protein gene *ACP-20* (adult-specific cuticular protein 20) had the highest expression (Table S3). At the same time point, the number of down-regulated genes was similar with that of up-regulated genes. Most proteins expressed by these genes were related to cellular immunity and apoptosis, including antimicrobial peptides (AAEL000625-PA, cecropin A precursor and defensin-C), toll-like receptor 13, cell death abnormality protein 1 isoform X2 and putative lysozyme-like protein. Moreover, three odorant-binding protein genes (*gene-LOC5577119, gene-LOC5571968* and *gene-LOC5567749*) and three serine protease easter genes (*gene-LOC5579410, gene-LOC5568135* and *gene-LOC5569890*) had a significantly down expression (Table S3). The top four proteins expressed by down-regulated genes were fizzy-related protein homolog involved in the pathway protein ubiquitination, AAEL000625-PA, angiopoietin-2-like (a growth factor which’s activities is mediated through the tyrosine kinase receptors [28]) and cecropin A precursor (Fig. 4A). Among the down-regulated genes at this time point, the *defensin-C* gene had the highest expression (Table S3).

Nearly half of up-regulated genes were undefined genes in the mosquitoes infected by intrathoracic inoculation at 7 dpi (Table S4). The top three proteins expressed by up-regulated genes were all the uncharacterized proteins (Fig. 4B). Among the down-regulated genes at this time point, the *gene-LOC110676601* had a higher expression, which could encode the protein VDAC (Voltage-dependent anion-selective channels, a pore-forming protein located in the outer mitochondrial membrane) (Table S4). Among the four genes with the highest down-regulated expression levels, only the protein expressed by *gene-LOC110680747* was a known protein, zinc finger protein 827-like, the other three were unknown (Fig. 4B). Moreover, five zinc finger protein genes (down-regulated: *gene-LOC110680747, gene-LOC5571062, gene-LOC5576563* and *gene-LOC110680309* and up-regulated: gene-LOC5573104) had differential expressions (Table S4). The zinc finger proteins are one of the most abundant groups of proteins and have a wide range of molecular functions. Recently it is reported that the zinc-finger protein ZFP36L1 can enhance host anti-viral defense against influenza A virus [29]. In addition, expressions of four venom allergen 5 genes (*gene-LOC5575393, gene-LOC5575399, gene-LOC5575400* and *gene-LOC5575401*) were significantly down-regulated. Previous findings showed that the venom allergen 5 has the association with deltamethrin resistance in mosquitoes [30], and *Ae. aegypti* venom allergen-1 promotes DENV and ZIKV transmission by activating autophagy in host immune cells [31]. Statistical comparisons of differentially expressed genes between the mosquitoes infected by intrathoracic inoculation at 2 dpi and at 7 dpi demonstrated that 3 genes exhibited a significantly different, including *defensin-C, chitinase 10* (*gene-LOC5570579*) and *gene-LOC5569955* (Fig.4 C).

### 3.7 Immune-related and metabolism-related process were found in EBIV-infected *Ae. aegypti*

KEGG pathway and GO enrichment analyses were performed to elucidate the biological functions and pathways in the EBIV-infected *Ae. aegypti* at 2 and 7 dpi. KEGG pathway results showed that genes were significantly enriched in four pathways at 2 dpi, including Toll and Imd signaling pathway, and endocytosis (Fig. 4D). However, the enrichment pathways were mainly focusing on biosynthesis and metabolism, including steroid hormone biosynthesis, N-glycan biosynthesis, ribosome, RNA degradation, and steroid biosynthesis at 7 dpi (Fig. 4E). Likewise, GO analysis results showed that immune-related process at 2 dpi and metabolism-related process at 7 dpi were significantly enriched (Fig. 4F and 4G).

## 4. Discussion

Herein, we evaluated the vector competence for EBIV and transcriptional response of *Ae. aegypti* to EBIV infection to provide the risk assessment of its transmission and its potential underneath mechanism. Our results demonstrated that *Ae. Aegypti* was susceptible to EBIV both trough oral feeding and intrathoracic-injection infection. The highest transmission rates in intrathoracic-injection mosquitoes could reach 90%, when compared to the oral feeding mosquito with 11.8%, which indicated the risk of EBIV transmission will greatly increase once the virus acquire much stronger ability to break through the midgut barrier. Compared to the vector competences of other viruses, the vector competence of *Ae. aegypti* for EBIV was not low and even better than other orthobunyaviruses, such as NRIV and BATV (Table S5). Due to the high infection rates in the ovary, there was a possibility of vertical transmission of EBIV. Transcriptome analysis results demonstrated that EBIV can alter the expression of mosquito genes related to immune function and metabolism-related process. It was worth noting that *defensin-C* and *chitinase 10* had been continuously down-regulated in the mosquitoes infected by intrathoracic inoculation.

Notably, after oral-infection, a few EBIV-positive saliva samples were appeared in the infected mosquitoes of which the viral loads in guts were more than 10^4^ PFU ml^−1^ (Fig. 1G), suggesting that EBIV might enter saliva when virus titer in the guts reached to 10^4^ PFU ml^−1^. When the midgut barriers were bypassed, EBIV was able to rapidly invade the whole mosquito bodies, including saliva and ovaries (Fig. 2G to 2H). These results indicated that *Ae. aegypti* may became an important vector to transmit EBIV to other vertebrates, and the midgut may be the main barrier to virus transmission [32]. In addition, it is not uncommon for orthobunyaviruses to spread themselves in immature life stages of mosquitoes through the mode of vertical transmission [33,34]. The highly EBIV infection rates in ovaries indicated the possibility of its vertical transmission (Fig. 1H and 2H).

As mentioned above, more than 90% of adult BALB/c mice succumbed to death when infected intraperitoneally with an extremely low dose of 1 PFU EBIV [20]. In addition, EBIV is considered more pathogenic for the adult mice model, compared to other viruses of the genus *Orthobunyavirus*. Adult mice generally show resistance to other orthobunyaviruses infections, while 3 weeks old or younger mice are more susceptible [20,35–37]. Our results showed that the mean viral titers of EBIV-positive saliva were more than 10^1.5^ PFU ml^−1^ (the average dose of EBIV per mosquito was more than 6.3 PFU) at 14 dpi in both oral-infected and intrathoracic virus inoculation mosquitoes, that is to say, which indicated a high risk of EBIV transmission to a vertebrate host by the mosquito. So far serological investigation of EBIV in vertebrate hosts has not been completed [19], but the dangers of EBIV for public health were fully reflected in animal models, which causes encephalitis with hepatic and immune system damages and a high mortality rate [20].

During the process of virus invasion, arboviruses must cope with innate immune responses and overcome several barriers of their mosquito vectors. In general, there are four major barriers, the midgut infection barrier (MIB), midgut escape barrier (MEB), salivary gland infection barrier (SGIB), and salivary gland escape barrier (SGEB) [38]. *Ae. aegypti* was moderately susceptible to EBIV after oral feeding, which 70.0% of mosquitoes with a midgut infection at 10 dpi, whereas the dissemination rate was only 38.1% at this time points, significantly lower than infection rate (*p* = 0.0432). Notably, the proportion of EBIV-positive saliva samples was 90.0% at 10 days post intrathoracic injection. Combined results of blood feeding and intrathoracic injection experiments in this study demonstrated that MEB might be the primary barrier for EBIV to systemically infect *Ae. aegypti*. An efficient MEB can result in that the virus replication is limited to the midgut or dissemination occurs inefficiently [39]. It is possible that the basal lamina of the midgut has effects on an effective dissemination [40]. Several studies have reported that there is an inverse relationship between LACV dissemination rates and thickness of the basal lamina of *Ae. triseriatus* [41,42]. Alternatively, the reason why EBIV has not an effective dissemination might be that EBIV is not able to overcome or evade the antiviral immune responses in midgut. Electron microscopy of midguts from mosquitoes infected via intrathoracic injection revealed the presence of a large number of autophagosomes and lysosomes, which suggests that autophagy may participate in EBIV infection. Recent studies about the effects of autophagy on arbovirus infection have demonstrated that autophagy may function as an antiviral defense [43–45]. Furthermore, it has been found that the replication of RVFV which belongs to the family *Phenuivridae* is limited by activation of autophagy in *Drosophila* [46]. However, there are some studies found that autophagy is benefit for flaviviruses replication and transmission in *Ae. aegypti* [47,48]. RNA interference (RNAi) pathway may also be the reason for inefficiently dissemination. RNAi, as one of innate antiviral responses, is able to limit the replication efficiency of Sindbis virus in *Ae. aegypti* and virus can replicate more rapidly and efficiently in RNAi pathway impaired mosquitoes [49]. Additionally, there are evidences suggesting that the competence for midgut infection efficiency is dose-dependent [38,50,51], which indicated that with the increasing viral titer in host’s blood, the threat of EBIV for public health will be more serious.

Because of the characteristics of arbovirus, which need to maintain their lifecycle by switching between vertebrate hosts and invertebrate vectors, there is a strong possibility that evolutionary adaptation will occur during this process [52,53]. For instance, a single amino acid substitution increased the susceptibility to *Ae. albopictus* and led to viruses to disseminate more rapidly from the midgut to secondary tissues [54]. The same situation also happens on ZIKV, the mutant enhances ZIKV infectivity in mosquitoes [55]. Therefore, once the virus evolves in nature to further adapt to the midgut barrier, there will be a great risk for public health, and it might be an emerging disease possibly influencing human or animal health and economy development.

In conclusion, the *Ae. aegypti* can be infected by EBIV and possibly transmit the virus, although the efficiency of dissemination and transmission is lower when compared to the large-scale epidemic mosquito-borne viruses, such as DENV and ZIKV. In future, further studies on vector competence for EBIV in other mosquito species are needed, and people should be aware of the potential spreading of EBIV by mosquitoes.

## Additional information

Table S1. Primer sequences of genes used for qRT-PCR

Table S2. The correlation between EBIV-induced cytopathic effect in BHK-21 cells and CT values of virus RNA by qRT-PCR.

Table S3. Differentially expressed genes in the mosquitoes infected by intrathoracic inoculation comparing to the mock-infected mosquitoes at 2 dpi.

Table S4. Differentially expressed genes in the mosquitoes infected by intrathoracic inoculation comparing to the mock-infected mosquitoes at 7 dpi.

Table S5. The vector competence of *Ae. aegypti* for several well-known MBVs.

## Acknowledgement

We acknowledge the assistance from staff of the Center for Instrumental Analysis and Metrology of Wuhan Institute of Virology, Chinese Academy of Sciences and the Analysis and Testing Center of Institute of Hydrobiology, Chinese Academy of Sciences. The work was funded by Alliance of International Science Organizations (No. ANSO-CR-PP-2020-05).

