## Supplemental Table 1,2 and 5 for "Vector competence and transcriptional response of *Aedes aegypti* for Ebinur Lake virus, a newly mosquito-borne orthobunyavirus"

**Supplements**

**Table S1. Primer sequences of genes used for qRT-PCR**

| Gene name | Primers |
| --- | --- |
| EBIV segment S | F：ATGGCATCACCTGGGAAAG  R：TTCCAATGGCAAGTGGATAGAA |

**Table S2. The correlation between EBIV-induced cytopathic effect in BHK-21 cells and CT values of virus RNA by qRT-PCR.**

| **EBIV** | **Virus titer**  **(PFU/mL)** | **virus dose used to inoculate flask (PFU)** | **Days Post-Inoculation that CPE Appeared** | **Ct values** |
| --- | --- | --- | --- | --- |
| No dilution | 1.3 x 10^6^ | 1.3 x 10^5^ | 2 | 17.35 |
| 10^-1^ | 1.3 x 10^5^ | 1.3 x 10^4^ | 2 | 20.7 |
| 10^-2^ | 1.3 x 10^4^ | 1.3 x 10^3^ | 2 | 24.0 |
| 10^-3^ | 1.3 x 10^3^ | 1.3 x 10^2^ | 2 | 27.5 |
| 10^-4^ | 1.3 x 10^2^ | 1.3 x 10^1^ | 3 | 30.6 |
| 10^-5^ | 1.3 x 10^1^ | 1.3 | 4 | 34.6 |
| 10^-6^ | 1.3 | 0.26 | - | 38.0 |
| 10^-6^ | 1.3 | 0.13 | - | 0 |
| 10^-7^ | 0.13 | 0.013 | - | 0 |

**Table S5. The vector competence of *Ae. aegypti* for several well-known MBVs.**

| **Virus name** | **Days post infection** | **Infection rate %** | **Dissemination rate %** | **Transmission rate %** | **References** |
| --- | --- | --- | --- | --- | --- |
| **Bunyamwera virus** | 7 | (31/103) 30.1 | (19/31) 61.3 | -- | [1] |
|  | 14 | (63/143) 44.1 | (51/63) 81.0 | 80.0 |  |
| **Ngari virus** | 7 | (2/58) 3.4 | 0.0 | -- | [1] |
|  | 14 | (4/96) 4.2 | 0.0 | -- |  |
| **La Crosse virus** | 5 | >30.0 | >70.0 | -- | [2] |
|  | 7 | >30.0 | >60.0 | -- |  |
|  | 9 | >50.0 | >80.0 | -- |  |
|  | 11 | >30.0 | >60.0 | -- |  |
| **Cache Valley virus** | 7 | (10/66) 15.2 | (4/4) 100.0 | -- | [3] |
|  | 14 | (9/82) 11.0 | (5/5) 100.0 | (3/10) 30.0 |  |
| **Batai virus** | 7 | (4/16) 25.0 | 0.0 | 0.0 | [4] |
|  | 14 | (3/44) 6.8 | 0.0 | 0.0 |  |
| **Zika virus** | 4 | >80.0 | -- | >80.0 | [5] |
|  | 8 | >90.0 | -- | >90.0 |  |
|  | 10 | 100.0 | -- | >90.0 |  |
| **Dengue virus-1** | 7 | (22/29) 75.9 | (20/22) 90.9 | (0/20) 0.0 | [6] |
|  | 14 | (35/47) 74.5 | (31/35) 88.6 | (16/31) 51.6 |  |
| **Dengue virus-2** | 7 | (16/53) 30.2 | (2/16) 12.5 | (0/2) 0.0 | [6] |
|  | 14 | (12/50) 24.0 | (2/12) 16.7 | (0/2) 0.0 |  |
| **chikungunya virus** | 3 | (22/23) 95.7 | (11/22) 50.0 | (1/11) 9.1 | [7] |
|  | 7 | (19/20) 95.0 | (14/19) 73.7 | (2/14) 14.3 |  |
|  | 14 | (20/21) 95.2 | (18/20) 90.0 | (3/18) 16.7 |  |
